# An efficient GUI-based clustering software for simulation and Bayesian cluster analysis of single-molecule localization microscopy data

**DOI:** 10.1101/2021.06.11.447933

**Authors:** Saskia Kutz, Ando C. Zehrer, Roman Svetlitckii, Gülce S. Gülcüler Balta, Lucrezia Galli, Susanne Kleber, Ana Martin-Villalba, Helge Ewers

## Abstract

Ligand binding of membrane proteins triggers many important cellular signaling events by the lateral aggregation of ligand-bound and other membrane proteins in the plane of the plasma membrane. This local clustering can lead to the co-enrichment of molecules that create an intracellular signal or bring sufficient amounts of activity together to shift an existing equilibrium towards the execution of a signaling event. In this way, clustering can serve as a cellular switch. The underlying uneven distribution and local enrichment of the signaling cluster's constituting membrane proteins can be used as a functional readout. This information is obtained by combining single-molecule fluorescence microscopy with cluster algorithms that can reliably and reproducibly distinguish clusters from fluctuations in the background noise to generate quantitative data on this complex process.

Cluster analysis of single-molecule fluorescence microscopy data has emerged as a proliferative field, and several algorithms and software solutions have been put forward. However, in most cases, such cluster algorithms require multiple analysis parameters to be defined by the user, which may lead to biased results. Furthermore, most cluster algorithms neglect the individual localization precision connected to every localized molecule, leading to imprecise results. Bayesian cluster analysis has been put forward to overcome these problems, but so far, it has entailed high computational cost, increasing runtime drastically. Finally, most software is challenging to use as they require advanced technical knowledge to operate.

Here we combined three advanced cluster algorithms with the Bayesian approach and parallelization in a user-friendly GUI and achieved up to an order of magnitude faster processing than for previous approaches. Our work will simplify access to a well-controlled analysis of clustering data generated by SMLM and significantly accelerate data processing. The inclusion of a simulation mode aids in the design of well-controlled experimental assays.

## 1 INTRODUCTION

Cells rely on transmembrane signaling to interact with the outside world. It is essential that cells can specifically and decisively be put into action in response to signals in a noisy and complex environment. To do so, mechanisms have evolved that allow the triggering of an all-or-none, lasting response if required. This often involves a threshold number of ligand-activated membrane molecules that recruit auxiliary molecules to form a larger assembly that, upon reaching threshold size, will switch the cell into a different state. These signaling assemblies appear as clusters of membrane proteins in the plasma membrane of cells. However, the clusters may represent only a small subfraction of the membrane protein in question in an otherwise randomly distributed larger population. Cluster algorithms can detect such active signaling clusters in a randomly distributed background if the exact spatial distribution of membrane proteins is known (Williamson et al., 2011; Khater et al., 2020). Cartography of membrane protein distribution at the nanoscale has been made possible by super-resolution microscopy approaches based on the sequential localization of single fluorescence-labeled proteins (SMLM, (Betzig et al., 2006; Rust et al., 2006; Heilemann et al., 2008)). Clustering has since developed into an essential readout for membrane protein function in many cellular processes. Over the last years, several cluster algorithms have been adapted specifically for the analysis of single-molecule fluorescence data of membrane proteins (Owen et al., 2010; Annibale et al., 2011a,b; Nicovich et al., 2017; Baumgart et al., 2019; Arnold et al., 2020; Pike et al., 2020). SMLM of membrane proteins and their cluster analysis still requires a high level of experimental and analytical expertise. To make cluster analysis more accessible, we here combined a selection of the latest clustering approaches with several useful computational features to speed up and streamline cluster analysis in a single, user-friendly software. Specifically, we implemented Bayesian Cluster Analysis, Ripley's-K-based clustering, DBSCAN (Rubin-Delanchy et al., 2015; Griffié et al., 2016), and ToMATo (Pike et al., 2020) for cluster analysis. We then compared the performance of these approaches on simulated and newly generated experimental data from different cellular systems. Furthermore, we implemented a pipeline for parallelized computing of cluster analysis and, as a result, could analyze even large datasets at a fraction of the time required before. Our software will simplify and accelerate cluster analysis as a readout of membrane protein function.

## 2 RESULTS

### 2.1 Structure of the GUI

To facilitate the use of parallelized Bayesian cluster analysis for the community, we developed an easy-to-use software called BaClAva (<u>Ba</u>yesian <u>Cl</u>uster <u>A</u>nalysis and <u>v</u>isualization <u>a</u>pplication) with a graphical user interface (GUI, Fig. 1). This software consists of a pipeline of three modules for simulations, clustering, and analysis that can be used independently via the GUI. The execution of thought experiments is essential in developing reliable experimental strategies and is especially important for data processing-intensive assays. To allow for the freehand design of ground-truth data while simulating realistic experimental output, we included a simulation module similar to FluoSim (Lagardère et al., 2020). This module allows the generation of user-defined clusters of molecules combined with a selected level of randomly placed background molecules. The results of this ground truth are then modeled as images resulting from an SMLM-experiment emulated based on experimental statistics of dye blinking, camera noise, and localization accuracy. The resulting image stack is localized using standard algorithms and can be used as an alternative to or alongside actual SMLM localization data in downstream clustering analysis. If desired, the generation of emulated microscopy images from the constructed localizations can be omitted, as exemplified in Fig. 3. This option is based on Griffié et al. (Griffié et al., 2016).

**Figure 1.**
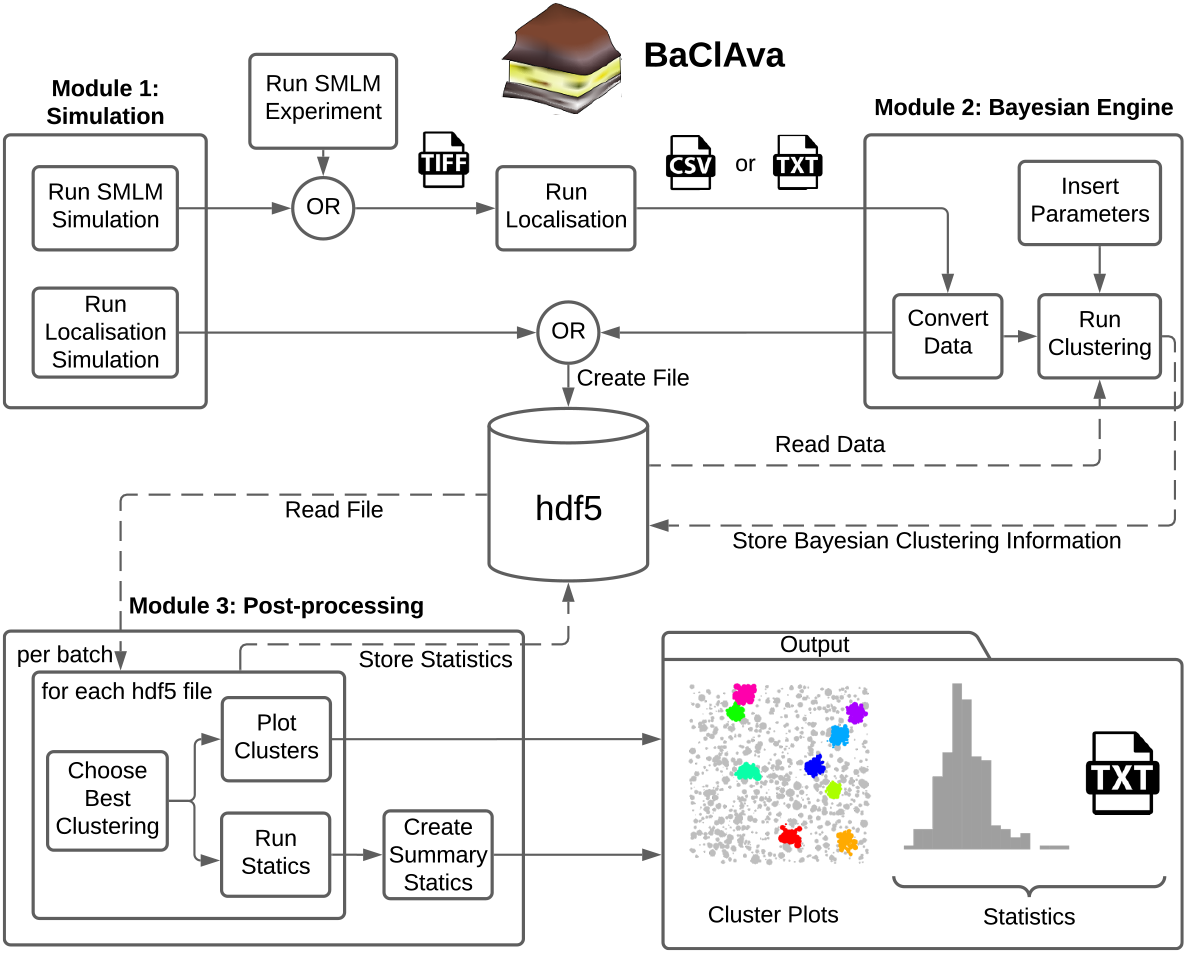
Overview of the software GUI. Schematic of the three independently usable software modes and organization of the software. Simulations can be prepared individually or as batches, and the localization results get exported as tiff or hdf5 files, depending on the simulation option. For the second module, the simulated data is imported from the hdf5 file, or experimental datasets can be imported in the form of a text or csv file. The user can set various parameters, most notably the cluster method, the type of computation, and additional Bayesian clustering parameters. This module's output, namely the scores and the labels for all proposals, is stored in the hdf5 file. In the final processing step, the original localization table and the Bayesian clustering module's output are used to produce the best cluster plots and the corresponding (batch) statistics. The statistics are exported as text files as well as plots.

**Figure 2.**
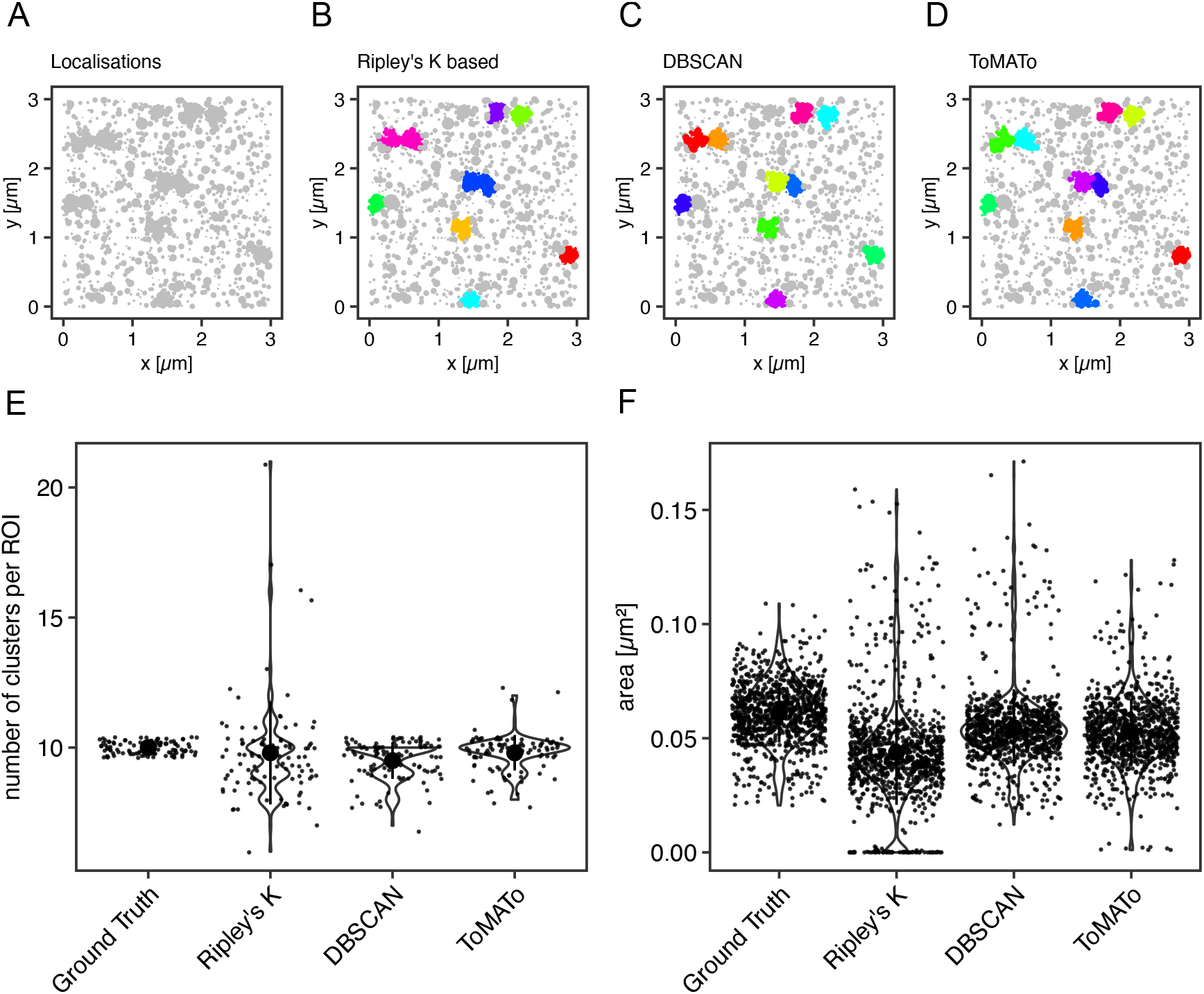
Comparison of cluster algorithms. **(A)** Example of one of 100 simulated ground truth datasets. **(B-C)** Cluster detection (colored) by **(B)** Ripley's-K-function-based implementation, **(C)** DBSCAN, and **(D)** ToMATo for this dataset demonstrating the respectively detected clusters. **(E)** Violin plots of the number of detected clusters in 100 simulations containing 10 ground-truth clusters for each of the algorithms implemented. The mean is emphasized as a black circle. Ten clusters were simulated, and the mean for Ripley's-K-based clustering was 9.8 +/− 2.0, for DBSCAN 9.5 +/− 0.7, and 9.8 +/− 0.7 for ToMATo. Note that the spread is significantly larger for Ripley's-K-based, DBSCAN never overcounted, and ToMATo was the most accurate overall. **(F)** Plot of all ground truth and recognized cluster areas. The ground truth data's cluster area has an average size of 0.061 +/− 0.013 μm^2^, the Ripley's-K-based clustering results in 0.044 +/− 0.023 μm^2^, DBSCAN in 0.055 +/− 0.017 μm^2^, and ToMATo clustering averages the area to 0.053 +/− 0.015 μm^2^ (mean +/− standard deviation).

**Figure 3.**
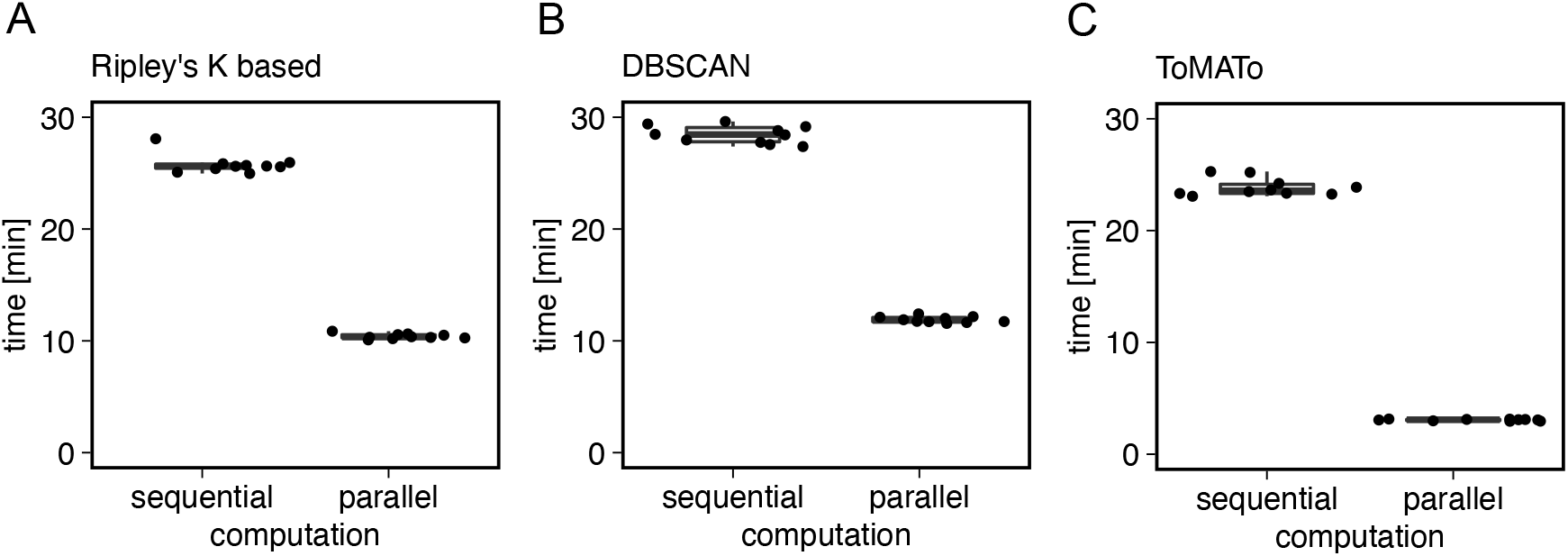
Computational costs for sequential and parallel implementations. Shown is the computational time of the Bayesian engine (in min) in sequential and parallel mode for **(A)** Ripley's-K-based clustering, **(B)** DBSCAN, and **(C)** ToMATo.

The second module is the clustering module, which analyzes single-molecule localization datasets in the format (X [nm], Y [nm], STDEV [nm]). Once the data are loaded into the software, the user can choose between ToMATo, Ripley's-K-based, or DBSCAN cluster analysis, define the desired parameter space for Bayesian analysis and select, whether the computation is done sequentially or in parallel. The third and final module allows the visualization and export of the results in a graphic or tabular form, including essential analytical parameters such as the number of clusters, cluster area, and cluster density.

To decrease the number of files stored on the computer disk, we decided to store all information in a Hierarchical Data Format (hdf5) (Fig. 1). The hdf5 format enables us to store the localization table (simulation or experimental), the Bayesian engine scores and labels, and further information in a single data file.

### 2.2 Benchmarking

First, we aimed to benchmark our cluster software on simulated clustering data. To do so, we generated 100 simulated images of clustered molecules, each containing ten clusters of 100 localizations. For example, see Figure 2A. These simulations were generated in the following way: Clusters were generated from single points ≥ 100 nm apart for each of which 100 localizations were generated by drawing from a normal distribution with a standard deviation of 50 nm. The random background was generated at a density of 111 localizations per μm^2^.

These data were then analyzed with the Bayesian model and the three different cluster detection algorithms. Figure 2 shows the simulated data and the corresponding clustering outputs. Since the cluster centers were set to be at least two standard deviations apart from each other, the individual clusters can be correctly identified by eye (Fig. 2A) and as well with DBSCAN (Fig. 2C) and ToMATo (Fig. 2D). In contrast and as shown before (Pike et al., 2020), the approach based on Ripley's K-function (Fig. 2B) fails to separate nearby clusters and thus commonly misidentifies cluster number and area (Fig. 2E, F). As previously shown, this behavior is due to the incapability of this approach to take into account the local density of the data points. In contrast, both DBSCAN and ToMATo could quantify both cluster number and overall cluster area quite accurately in the majority of simulations (Fig. 2E, F).

These methods in the Bayesian cluster approach rely not on a single set of parameters but instead on a continuum of so-called proposals, defined sets of values computed to cover an ample parameter space to find an overall optimum of cluster identification (Rubin-Delanchy et al., 2015; Griffié et al., 2016). While this approach has proven to lead to superior results, it is necessarily computationally costly. We aimed to overcome this problem to increase processing speed and thus experimental throughput.

So far, the cluster proposals' calculation in Bayesian analysis is done in nested for-loops on a single CPU core. Since the individual cluster proposals are independent of each other, the processing could also be implemented in parallel. This means that the program uses multiple CPU cores instead of a single core and therefore calculates multiple proposals at the same time. In our software, we implemented the parallelized computing of Bayesian cluster analysis and compared the results with the sequential computational approach.

We first used ten simulations to benchmark the clustering methods described above in Bayesian analysis. We found that typical runtimes for Ripley's K-based and DBSCAN clustering were 25.78 ± 0.86 min and 28.45 ± 0.78 min, respectively (mean ± standard deviation). The ToMATo implementation from the RSMLM package (Pike et al., 2020) had a runtime of 23.87 ± 0.80 min (mean ± standard deviation, Fig. 3). By parallelizing the clustering and scoring process to multiple cores, we found the computation time to decrease by 60% for Ripley's K-based, 10.41 ± 0.23 min, and DBSCAN, 11.90 ± 0.27 min (Fig. 3). For the ToMATo implementation, the computational time decreased by one order of magnitude to 3.062 ± 0.072 min. We concluded that parallelization significantly reduced processing time for Bayesian cluster analysis.

Next, we aimed to investigate several known sources of error in clustering single-molecule localization microscopy data. An important source of error in the cluster analysis of SMLM data is caused by multiple localizations of the same fluorescent molecule generated by most SMLM approaches that necessarily generate a cluster of localizations from every single fluorophore. Consequently, this fact must be considered for declaring any statement on fundamental information such as cluster size in terms of area and number of molecules in the cluster. By simulating blinking SMLM data with realistic blinking statistics, we determined how dense the underlying molecules must be for proper cluster detection. The simulations of (*d*)STORM experiments were generated in the following way: Cluster areas were generated by randomly distributing 40 non-overlapping clusters with an area of 0.0078 μm^2^ (diameter = 50 nm). Their molecular density was increased from 0.71 ± 0.25 × 10^3^ μm^−2^ to 6.24 ± 0.63 × 10^3^ μm^−2^. The random background was generated at a density of 607 molecules per μm^2^ for sparse clusters up to a density of 328 molecules per μm^2^ for dense clusters. The blinking parameters were k*_on_* = 0.01 s^−1^ and k*_off_* = 10 s^−1^. The FWHM of the PSF was set to 200 nm with an intensity of 2007. The pixel size of the camera was set to 0.096 μm, which is identical to the pixel size of the Evolve Delta 512 Photometrics camera on our microscope. The exposure time was set to 10 ms, which is the exposure time we use in experiments with Alexa Fluor 647 dyes, and as in a (*d*)STORM experiment, 50,000 frames were acquired. The localization procedure and grouping were done in SMAP (Ries, 2020). The obtained localization table was used for the Bayesian Analysis. The results are visualized in Figure 4.

**Figure 4.**
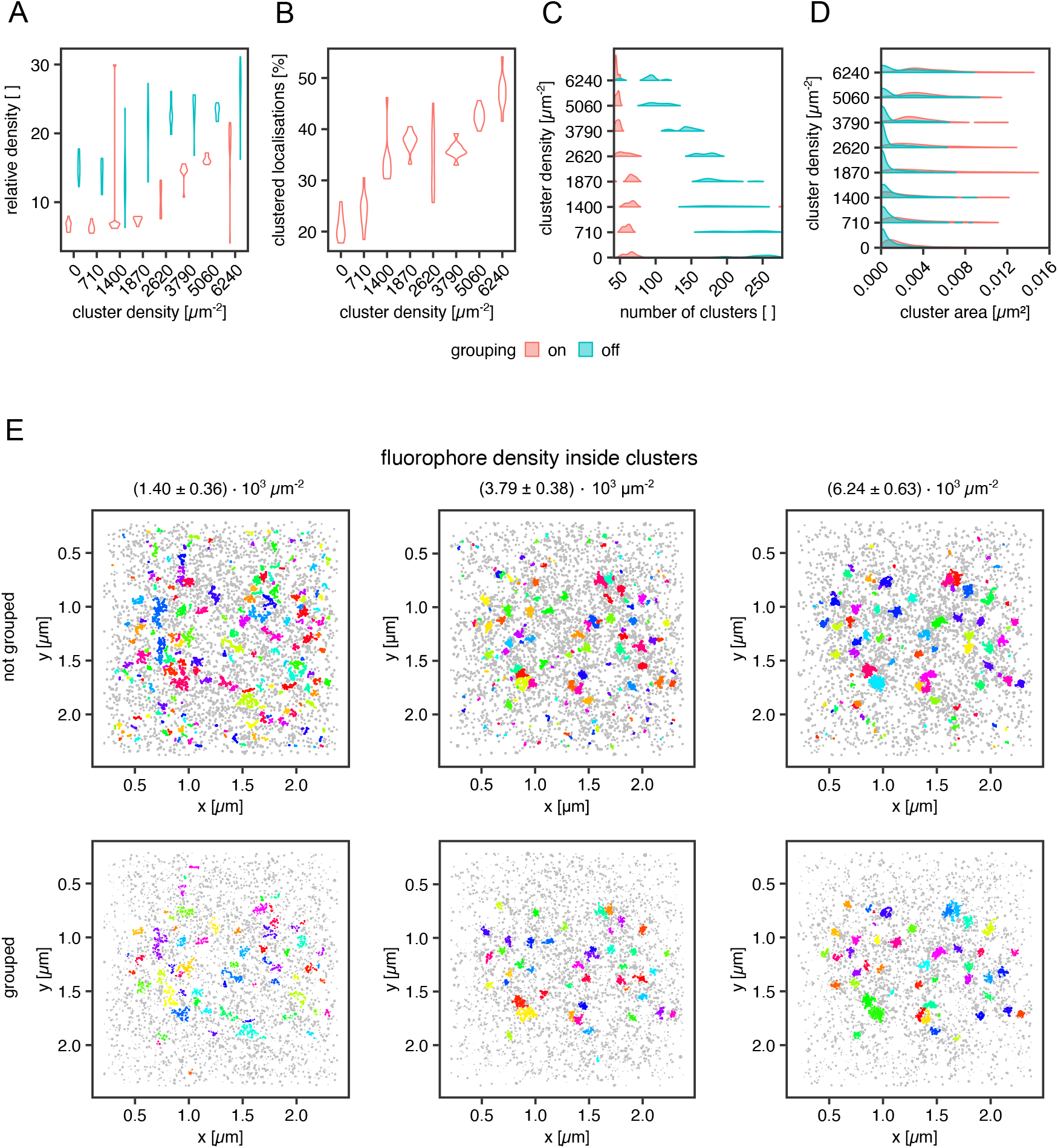
Influence of fluorophore blinking on clustering. **(A)** Violin plot for the relative density of the clusters vs. the background with and without grouping applied, **(B)** Violin plot of the percentage of the clustered localization with grouping, **(C)** Violin plot of the number of clusters per ROI with and without grouping applied, **(D)** Violin plot of the areas of the clusters with and without grouping, **(E)** Examples for clustering of a random distribution of fluorophores and 40 clusters at a density of 1.40 +/− 0.36 × 10^3^ μm^−2^ (left column), 3.79 +/− 0.38 × 10^3^ μm^−2^ (middle column) and 6.24 +/− 0.63 × 10^3^ μm^−2^ (right column). Each dataset was analyzed in SMAP with and without grouping. The cluster analysis was performed by the Bayesian engine plus ToMATo.

In Figure 4A, the cluster to background density for grouped and non-grouped data is shown. For both cases, the relative density increases with increasing cluster density and a smaller spread of the distributions for grouped data, whereas the non-grouped data distributions show a broader spread, indicating the efficiency of the grouping function in SMAP. Additionally, the number of clusters per region of interest (ROI) is reduced by the grouping, which removes clusters caused by single blinking fluorophores (Fig 4C, E). The higher relative density of fluorophores in clusters compared with background localizations indicates that a local density threshold must be surpassed to render the interpretation of cluster data independent of fluorophore blinking properties. As shown in Figure 4C, the number of clusters is constant for grouped data up to a concentration of 2.62 ± 0.39 × 10^3^ μm^−2^ localizations. For higher concentrations, the number of clusters approaches the ground truth of 40 clusters. Without grouping, the number of identified clusters decreases with increasing fluorophore concentration, reflecting a higher relative enrichment of fluorophores inside the clusters than outside them. The improved situation for grouped data is also visible in Fig 4B, showing that the percentage of clustered localizations increases with increasing fluorophore density. For the best cluster result in these simulations, more than 30% of the localizations must occur in clusters, and a relative density (localization density inside vs. outside of cluster) threshold of 10 must be overcome for the localizations inside clusters versus outside.

Moreover, the cluster size (Figure 4D), meaning the area covered by localizations in a cluster, shows the influence of background localizations on the data distribution. Cluster area increases in size for grouped data starting from a concentration of 1.87 ± 0.39 × 10^3^ μm^−2^. For the non-grouped data, there is a significant proportion of very small clusters at all concentrations. This cluster population is not present for the grouped data, indicating that these clusters emerge from multiple detections of a single fluorophore, i.e., blinking. For a density of 2.62 ± 0.39 × 10^3^ μm^−2^ molecules and higher, a second population emerges in the non-grouped data distributions, which corresponds to the main population in the grouped distributions. Therefore, they can be considered correctly identified clusters. Similarly, from 2.62 ± 0.39 × 10^3^ μm^−2^ molecules onwards, the number of clusters per ROI decreases. As demonstrated in Figure 4E, small background clusters are removed with the grouping functionality (top row vs. bottom row) and with increasing fluorophore density within the clusters (from left to right). As expected, the ground truth clusters become more apparent when the number of clustered molecules is increased even in the non-grouped data, indicating that single fluorophore blinking has a significantly reduced impact on density-based cluster identification for denser clusters. We concluded that grouping is essential in the detection of smaller clusters.

Finally, we aimed to apply our algorithm to experimental data from single-molecule localization experiments of intact cells. We used standard controls in the field for non-clustered and clustered molecules respectively at the plasma membrane. The lipid-anchored glycosylphosphatidylinositol-coupled green fluorescent protein GPI-GFP should be more or less homogeneously distributed and functioned as the negative control. The clathrin-light chain (CLC), of which dozens of copies are incorporated into every ~ 150 nm diameter clathrin-coated pit and thus appears strongly clustered, served as the positive control. In order to keep our results comparable, all molecules of interest were tagged with a GFP protein, and the (*d*)STORM dye Alexa Fluor 647 was bound to the GFP via anti-GFP nanobodies in all experiments (Ries et al., 2012). From the simulation work, we know that the cluster results for GPI-GFP should show a wide range of cluster areas, whereas, for the CLC, we expect to yield well-defined cluster areas. Finally, we asked whether we could detect clustering for the transmembrane receptor CD95, as the receptor activation via its ligand may trigger apoptosis or tumorigenesis of cancer cells and has been suggested to result in the formation of high order molecular clustering (Martin-Villalba et al., 2013). CD95 was likewise labeled via GFP and AF647 nanobodies.

The reconstructed images in Fig. 5 of these three proteins show differences in the spatial distribution of the localizations. For GPI-GFP imaged in CV-1 cells in Fig. 5A, the localizations are evenly distributed, and the cluster maps for the zoom-ins show small clusters, which are probably due to the blinking of the Alexa Fluor 647 dye. In contrast, in Fig. 5B, the CLC imaged in HeLa cells show well-defined clusters in agreement with clathrin-coated pit size (Fig. S1) with little background localizations, as seen in the cluster maps of the zoom-ins. The CD95 receptor in T98G glioblastoma cells presents a localization distribution with smaller clusters and more background localizations than CLC. The cumulative distribution of the cluster areas of several cells for each condition in Fig. 5D reveals that GPI and CLC exhibit distinct distributions of their respective cluster areas in agreement with expectations. The cumulative distribution of cluster areas for the CD95 receptor is positioned between the two controls, demonstrating that CD95 forms small clusters likely consisting of around 0.54 molecules/nm^2^ in the plane of the membrane.

**Figure 5.**
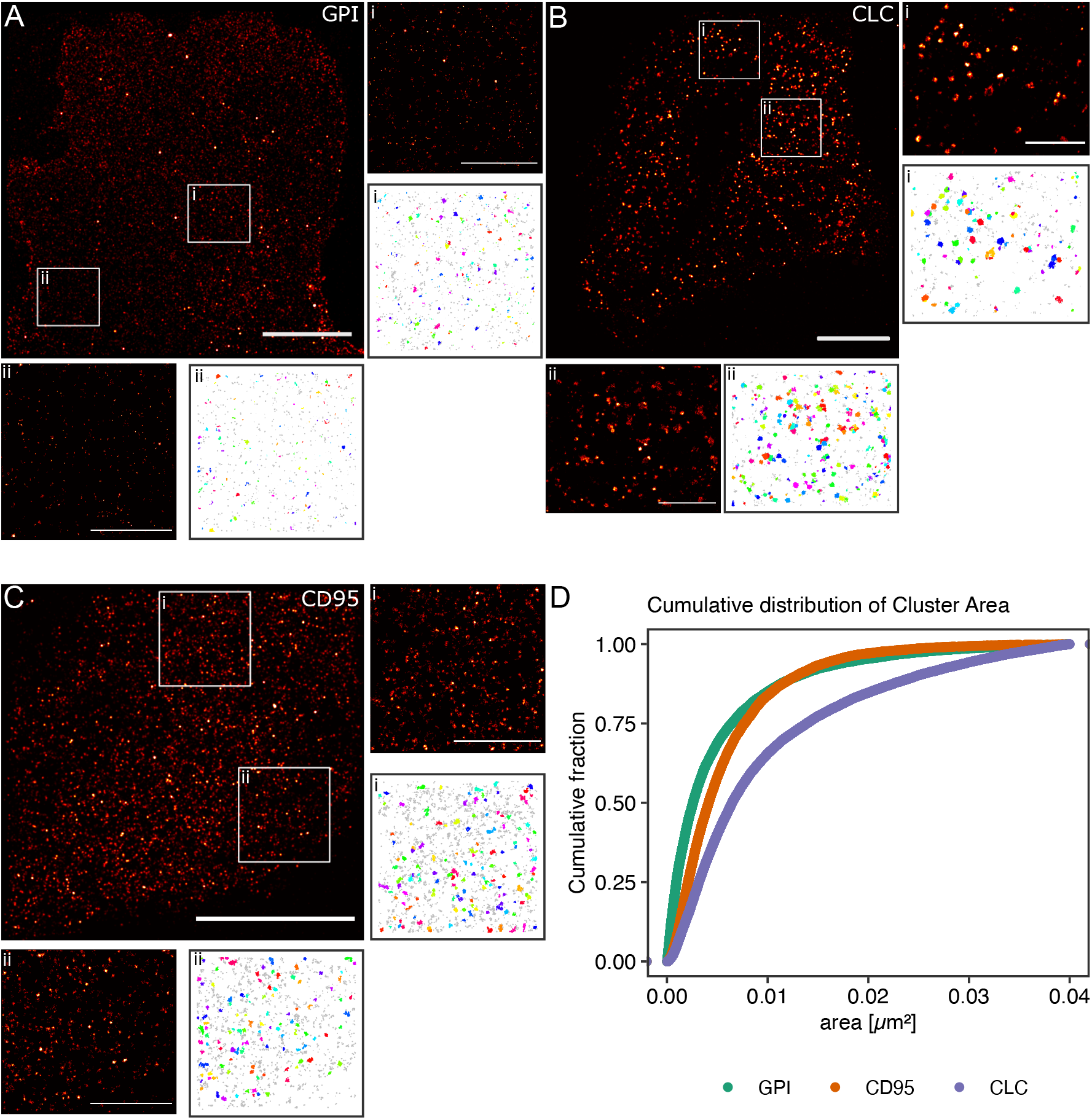
Practical application of the Bayesian engine on three different target molecules. Three different molecules were coupled to GFP and stained with nanobodies labeled with Alexa Fluor 647. **(A)** GPI-GFP in CV-1 cells, **(B)** CLC-GFP in HeLa cells, and **(C)** CD95-GFP in T98G cells. Scale bars are 10 μm for the large reconstructed images and 3 μm for the zoom-ins. **(D)** Plot of the cumulative distribution of cluster areas for the three target molecules.

## 3 DISCUSSION

Here we present a user-friendly software solution for cluster analysis of SMLM data. Our software significantly reduces processing time and allows the user to select different algorithms to identify and quantify cluster formation.

The simplest cluster algorithms, such as nearest-neighbor algorithms, can answer whether areas of above-average concentration, clusters, exist in a field of view (Endesfelder et al., 2014). For a more detailed analysis of clusters found in cellular membranes, Ripley's K-function can provide answers to the length scale of interparticle distances and the proportion of entities found in clusters in a given dataset (Owen et al., 2010). However, this function is prone to artifacts intrinsic to single-molecule fluorescence-based microscopy approaches, which lead to small local clusters due to the blinking behavior of individual fluorophores. To overcome errors due to blinking, approaches have been developed to determine the degree of clustering in challenging experimental circumstances, such as for dense membrane molecules by varying the dye density (Baumgart et al., 2019; Arnold et al., 2020).

To understand the functional underpinnings of cluster formation in cell biology, a qualitative view of clustering is not sufficient, but reproducible, robust quantitative assays are required. One of the first ideas put forward was to use Ripley's function not on the entire sample but on individual localizations convoluted with a search radius and a clustering threshold for cluster identification in dense backgrounds. To facilitate the differentiation of Ripley's K function on the entire sample or individual localizations, we termed the second case Ripley's-K-based clustering.

As shown before (Pike et al., 2020), Ripley's K-based clustering cannot adequately determine clusters in samples with variations in cluster density and final cluster size. Therefore, density-based clustering methods, like DBSCAN (density-based spatial clustering of applications with noise), have been adapted for SMLM data. Here, the search parameters are the search radius and the minimal number of data points within this search radius. Points for which these parameters are valid are counted towards this cluster. After identifying all clusters, the remaining data points are assigned to the background, and other parameters such as the individual cluster area and density can be extracted from the data.

Even though density-based methods, like DBSCAN, can handle datasets with density variations, they still often fail to separate individual clusters that are very close to one another but are easily distinguishable by eye. Additionally, it has been shown that the detection of clusters by DBSCAN and Ripley's-K-based clustering (Pike et al., 2020) can be sensitive to even small changes in the analytical parameters, possibly leading to artifactual results. One of the latest introductions to SMLM cluster analysis is persistence-based clustering which is based on density estimation. The introduced scheme is called Topological Model Analysis Tool (ToMATo, (Pike et al., 2020; Chazal et al., 2012, 2013)). In contrast to the above-mentioned density-based methods, in this algorithm, local maxima in molecular density are identified and termed clusters by introducing a density gradient generated by creating a path emanating from a molecule to its neighbors and using the intermolecular distance as a measure of density. If this intermolecular distance increases, the border of a cluster may be reached. Such a local maximum in distance or minimum in density may be a saddle point between clusters or define the outer perimeter of a cluster. A threshold value defines the persistence of a cluster from its center into space. Clusters with persistence smaller than the threshold are assigned to neighboring clusters, or they are deemed background. As a result, ToMATo allows for a separation of even partially overlapping clusters, and additionally, the output clustering results are less sensitive to analytical input parameters as compared to Ripley's K-based clustering and DBSCAN. Further clustering methods for SMLM data are based on Voronoï tessellation (Levet et al., 2015), which detects clusters based on polygonal regions. Voronoï tessellation intrinsically generates contours of regions of density that may also be used for boundary detection of cells. Recently, machine-learning has been employed to improve cluster detection (Williamson et al., 2020), but the number of input neurons limits the correct processing of the underlying information. The Bayesian engine's main drawback is that due to the calculation and scoring of thousands of cluster proposals for optimal results, the process is significantly slowed down compared to traditional methods with a single cluster proposal, hampering the routine use of this method. On the other hand, ToMATo clustering parameters are determined based on a persistence diagram which can cause user bias.

To overcome these limitations and provide accessible GUI-based software for state-of-the-art cluster analysis, we implemented Ripleys-K-based clustering, DBSCAN, and ToMaTo in a common software that allowed for parallel computing. In this software, we first improved the Bayesian engine's speed by implementing parallel computation and introduced ToMATo clustering to the Bayesian engine, thereby dramatically decreasing computational time. In combination with the software GUI, the Bayesian engine has an improved user experience and processing speed, which we hope will make state-of-the-art SMLM cluster analysis available in many laboratories.

During an SMLM measurement of several thousands of frames, a fluorescent molecule may cycle several times between a bright and a dark state, and thus, one molecule may be detected multiple times within a radius determined by its localization precision. As a result, it is impossible to differentiate between a single molecule detected several times in different frames and different molecules in close proximity detected in different frames. This is, of course, especially problematic in cluster analysis, where localizations are processed first without bias. To overcome this problem, it is important to develop an understanding of the degree of influence of blinking in the dataset at hand. As we showed in Fig. 4, the number of the small clusters resulting from dye blinking decreases with increasing molecular density within the clusters while keeping the actual cluster size constant. Thus, there is an intrinsic threshold for relative localization densities inside and outside clusters that render blinking irrelevant. This holds true under the assumption that all localizations are caused by dyes bound to molecules of interest, and no false localizations are present in the sample. Below this threshold, the number of detected clusters is highly overestimated, and the cluster radii are dramatically underestimated. From simulations, we know that such single-molecule clusters can be detected as sub-peaks within clusters at low-density ratios. Increasing the density ratio now increases the chance that clusters are quantified at their true size. It is common in SMLM data analysis that multiple temporally and spatially closely correlated localizations are grouped together in a final reconstruction and are thus counted as a single molecule. In clustering, this procedure reduces the number of small background clusters dramatically, and we analyze this effect in depth in Figure 4. Our grouping is based on the blinking behavior of the most used (*d*)STORM dye Alexa Fluor 647, which we also used in our experiments. Likewise, we also based our simulations on the blinking behavior of this dye (Heilemann et al., 2008; Dempsey et al., 2011). In order to detect clusters of smaller density ratios and smaller sizes either or both, the cluster detection may be improved by changing the dye or even the SMLM method from (*d*)STORM to DNA-PAINT as shown in (Jayasinghe et al., 2018).

Microscopy experiments in cells are much more complex than the corresponding *in-silico* experiments because many different known and unknown cellular processes are involved in the temporal and spatial organization of the cell molecules and may interfere in the process studied. Therefore, we chose highly abundant molecules as cellular controls for clustering experiments. A simple positive control is clathrin-coated pits expressed at a well-defined radius of around 80 nm (A ≈ 0.02 μm^2^) (Sochacki et al., 2017). Negative controls for clustering in a cellular environment are far more challenging to identify because natural cellular signaling processes result in a spatial and temporal reorganization of the involved molecules, and many membrane molecules exhibit clustering of some extent (Gowrishankar et al., 2012; Saka et al., 2014; Baumgart et al., 2016; Kalappurakkal et al., 2020). Therefore, the influence for altering the negative control's organization by cell processes should be kept at a minimum, and an artificially introduced protein that is only anchored to the outer membrane of the plasma membrane and has no natural interaction partners, such as GPI, is the ideal option (Li et al., 2020). These extreme cases of clustering and non-clustering probes can be well differentiated in their reconstructed images as well as their cumulative distribution functions. Proteins with so far unknown spatial distribution on the plasma membrane, such as the transmembrane receptor CD95, should present a behavior between these two extremes. If they are less clustered, they should tend towards a behavior similar to GPI, and with increasing cluster areas, they should tend towards a distribution similar to clathrin-coated pits. Since CD95 can be found at the plasma membrane as monomers or homodimers and homotrimers (Micheau et al., 2020), it should be detected as small clusters, as observed in Figure 5. We conclude that our software can correctly distinguish between unclustered molecules and clusters of even small size and a few molecules in number.

Taken together, our work allows the implementation of single-molecule clustering analysis at a high rate of data throughput for beginning users. We expect our work to accelerate research in this area significantly and to contribute to the acceptance of reproducible standards in clustering data analysis. In future work, other analytical methods such as Voronoï tessellation (Andronov et al., 2018; Levet et al., 2015) and extensions to 3D (Griffié et al., 2017) and dual-color co-clustering (Jayasinghe et al., 2018) may be implemented, and the processing speed may be further improved, i.e., by the implementation of GPU-processing.

## 4 MATERIAL AND METHODS

### 4.1 Cell culture and preparation

CV-1 cells were cultured in a standard DMEM medium (1X, Gibco) supplemented with 10% FBS (ThermoFisher) and 1% GlutaMax (100X, Gibco by Life Technologies). Stable HeLa CLC-GFP cells were cultured in the same medium with an additional 1% Penicillin-Streptomycin (Sigma), and for the T98G CD95-GFP cells, 1% sodium pyruvate (stock: 100mM, Gibco) was added to the medium. The vector CD95-GFP was infected into the cells with a lentiviral construct. The cells were then FACS sorted for the stably transfected clones. All cell lines were regularly tested for mycoplasmas and only used when tested negative. For the seeding of the cells, 18 mm diameter #1.5 glass slides (VWR) were cleaned in an ultrasound bath for 20 min using 2% Hellmanex III (Hellma) and 70% ethanol, respectively. Afterward, the glasses were dried and plasma cleaned for another 30 min.

### 4.2 Cell staining

#### CV-1 GPI-GFP cells

Transfection of GPI-GFP into CV-1 cells was done with lipofectamine 3000 following the standard protocol (lipofectamine protocol by Invitrogen/ThermoFischer). Cells were treated with trypsin-EDTA and seeded on the glass slides for incubation of 24 h (densities: 6 × 10^6^ cells/ml for CV-1, 7 × 10^4^ cells/ml for HeLa CLC-GFP and T98G CD95-GFP). The transfected CV-1 cells were fixed with prewarmed 4% PFA with 0.2% GA in PBS for 20 min at 37 °C. Then, cells were quenched with freshly prepared 0.1% NaBH_4_ in PBS for 7 min at room temperature and extensively washed. Cells were blocked in two steps: for 30 min with ImageIT, followed by 4% goat serum in 1% BSA in PBS for 1 hr. CV-1 GPI-GFP cells were stained with anti-GFP nanobodies (FluoTag-Q anti-GFP) labeled 1:1 with Alexa Fluor 647 from NanoTag Biotechnologies GmbH at a concentration of 50 nM for 1 hr. Afterward, cells were postfixed with 4% PFA and 0.2% GA in PBS for 20 min and quenched with 0.1% NaBH_4_ in PBS for 5 min at room temperature.

#### HeLa CLC-GFP cells

HeLa CLC-GFP cells were fixed with prewarmed 4% PFA in PEM for 20 min at 37 °C and quenched with NH_4_Cl in PBS for 5 min at room temperature. After quenching for 5 min with 0.2% saponin in PEM, the cells were blocked with 4% goat serum in 1% BSA in PEM for 1 h. HeLa CLC-GFP cells were stained with the NanoTag Biotechnologies GmbH nanobody for 30 min at a concentration of 50 nM and afterward post-fixated with 4% PFA in PEM for 20 min at room temperature. The cells were quenched with NH_4_Cl in PBS for 5 min. In between all steps, the HeLa cells were extensively washed with PEM

#### T98G CD95-GFP cells

The T98G CD95-GFP cells were fixed for 20 min at 37 °C with prewarmed 4% PFA plus 0.2% GA in PEM and quenched with freshly prepared 0.1% NaBH_4_ in PEM for 7 min. T98G cells were permeabilized with 0.2% saponin in PEM for 5 min and blocked with 4% goat serum in 1% BSA/PEM for 1 hr. The cells were stained with the NanoTag Biotechnologies GmbH nanobody at a concentration of 50 nM for 30 min and post-fixated with 4% PFA with 0.2% GA in PEM for 20 min at room temperature. For the post-quenching, the cells were incubated in 0.1% NaBH_4_ in PEM for 7 min. In between all steps, the cells were extensively washed with PEM.

### 4.3 (*d*)STORM imaging

The fixed and stained samples were mounted and imaged in beta-mercaptoethanol and GLOX (2.5 mg/ml glucose oxidase, 0.2 mg/ml catalase, 200 mM Tris-HCl pH 8.0, 50% glycerol) as imaging buffer (10:1). The (*d*)STORM images were acquired on a home build TIRF microscope as described in (Albrecht et al., 2016). For the imaging, an Olympus 60x, 1.49 NA back focal plane TIRF objective was used to reach a pixel size of 96 nm. The samples were illuminated with a 639 nm laser (Changchun New Industries Optoelectronics Tech. Co., Ltd.) at powers of 0.008 - 0.015 mW/μm^2^. For the acquisition of the (*d*)STORM images, a water-cooled and back-illuminated Photometrics EMCCD camera with 512 × 512 pixels at a pixel size of 16 × 16 μm was used for the acquisition of 30,000 frames at an exposure time of 10 ms. The EMCCD camera was calibrated before the data acquisition, and the image acquisition was controlled with MicroManager.

### 4.4 (*d*)STROM reconstruction

The acquired and simulated (*d*)STORM datasets were localized using SMAP (Ries, 2020). Important camera and acquisition parameters were extracted from the metadata file, which had been saved with the data. Furthermore, the electron multiplier (EM) gain was set to 300, and the conversion factor to 6.7 (analog to digital units to photons). The minimum distance between two candidate peaks in order to be fitted separately was set to 7 pixels. For the point-spread function (PSF) fitting, the following parameters were set to a differential of Gauss with sigma = 1.2, dynamic factor = 1.7, and free PSF, using the workflow 'set Cam parameters’.

### 4.5 Grouping in SMAP

The grouping procedure is a part of SMAP, which we used for the reconstruction. The number of frames, dT, for which a single molecule can be non-fluorescent but still be grouped with the first localization of that molecule was set to dT = 1. The distance, dX, the centroid of a single molecule can be shifted in the image plane between two consecutive frames, but still, be grouped with the first localization of the molecule, was set to dX = 1. These are the standard values in SMAP, and they were identified as the optimal parameter values for out (*d*)STORM experiments.

### 4.6 Simulations

#### Simulations used for cluster algorithm comparison

The simulations were done with an adapted simulation code published by Rubin-Delanchy et al. (2015). The number of clusters, the number of molecules inside each cluster, the corresponding standard distribution for the cluster size, and the background percentage were set depending on the analysis. Unlike the original publication, the cluster centers are set to be at least two standard distributions apart from each other. In total, 100 simulations were done for each case.

#### Simulations used for computational time evaluations

Ten simulations with a standard deviation of 50 nm, 10 clusters with 100 molecules each, and 50% of the total number of localizations in the background were used to determine the computational cost for the three cluster algorithms combined with the Bayesian engine.

#### Simulations with blinking molecules

Simulations were prepared in Fluosim (Lagardère et al., 2020). For the simulation of the sample staining, a geometry file was created with a python script. The field of view had a size of 25 × 25 μm^2^ and was composed of 40 randomly distributed, non-overlapping clusters with a diameter of 50 nm. The clusters were positioned with a minimum distance of 500 nm from any border of the sample. The background image was an image of the Evolve 512 EMCCD camera (Photometrics) with a size of 26 × 26 μm^2^. The pixel size matched the pixel size of our experimental setup. Each pixel's noise values were not considered because only the pixel shape was used in the further course. The number of molecules was set to 4000 to match the density of optimal CV-1 GPI-GFP samples stained with anti-GFP nanobodies labeled with Alexa Fluor 647. For a fixed period (5 s – 50 s), the molecules were diffusing within the field of view with a coefficient of 0,01 μm^2^/s. A binding rate of 0,997-1,007 s^−1^ was set to allow cluster formation inside the clusters. Outside the designated cluster areas, the binding rate was set to 0 s^−1^. After the binding period, the molecules were freely diffusing for 50 s. During this time, the binding and unbinding rates within the clusters were set to zero and set to 0,997 - 1,007 s^−1^ outside of the cluster areas, thereby causing a homogeneous distribution of background molecules.

For simulating an actual SMLM experiment, the fluorophores' blinking parameters and the optical properties of the fluorescence emission were set accordingly. The on-rate was 0.01 s^−1^, and the off-rate 10 s^−1^, based on an estimated 1:1000 ratio in an SMLM experiment. For fitting the point-spread function, full-width at half maximum was fixed at 200 nm with a fluorescence emission intensity of 2007. As in a microscopy experiment, 5000 frames were acquired of the simulated sample, and the exposure time was set to 10 ms/frame. The output tiff-file was localized in SMAP with the standard parameters used for SMLM imaging. The camera parameters were the default values of the Delta 512 as given by its metadata file.

### 4.7 Computational runtime measurements

To evaluate the implemented cluster algorithms' speed, we used a standard 64-bit laptop computer running Linux (Ubuntu 18.04.5 LTS), equipped with GNOME 3.28.2, 7.7 GiB of memory, and 4 Intel® Core™ i5-6200U CPU @ 2.30 GHz processors. The R library ‘tictoc’ (Izrailev, 2014) was used to measure the time needed for each dataset to be processed.

### 4.8 Bayesian analysis

#### Cluster algorithms

The Ripley's-K-based and DBSCAN cluster algorithms used were written by (Rubin-Delanchy et al., 2015; Griffié et al., 2016). The code was adapted for improvement by using functions from several R packages and the ToMATo cluster algorithm for SMLM data adapted from the R package RSMLM (Pike et al., 2020). The library ‘doParallel’ was used for parallel implementation (Analytics and Weston, 2014).

#### Bayesian parameters

All Bayesian cluster scorings were done with the same set of parameters. The percentage of background localizations was set to 50%, and the Dirichlet process's concentration coefficient was 20. The optimal cluster parameters (radius and threshold) were searched in the sequences 5 to 300 for the first parameter and 5 to 500 for the second parameter in steps of 5.

#### Statistical analysis

The statistical comparison was performed with a self-developed R script.

## AUTHOR CONTRIBUTIONS

Funding acquisition: AMV, HE; Software: SK, RS; Sample preparation & acquisition: SK, ACZ; Simulations: ACZ, RS, SK; Writing – original draft: SK, HE; Resources: GSGB, LG, SK, AMV; Writing – review & editing: SK, ACZ, HE;

## ACKNOWLEDGMENTS

We would like to acknowledge the assistance of the Core Facility BioSupraMol supported by the DFG. The authors would like to thank the HPC Service of ZEDAT, Freie Universität Berlin, for computing time. We acknowledge support by the Open Access Publication Initiative of Freie Universität Berlin.

## DATA AVAILABILITY STATEMENT

The software package and the user guide are available at the Github repository “https://github.com/saskiakutz/BaClAva”. The GUI is written with the python package PyQT5, and the python - R connection was enabled rpy2.

